# An epigenetic biomarker of aging for lifespan and healthspan

**DOI:** 10.1101/276162

**Authors:** Morgan E. Levine, Ake T. Lu, Austin Quach, Brian H. Chen, Themistocles L. Assimes, Stefania Bandinelli, Lifang Hou, Andrea A. Baccarelli, James D. Stewart, Yun Li, Eric A. Whitsel, James G Wilson, Alex P Reiner, Abraham Aviv, Kurt Lohman, Yongmei Liu, Luigi Ferrucci, Steve Horvath

## Abstract

Identifying reliable biomarkers of aging is a major goal in geroscience. While the first generation of epigenetic biomarkers of aging were developed using chronological age as a surrogate for biological age, we hypothesized that incorporation of composite clinical measures of phenotypic age that capture differences in lifespan and healthspan may identify novel CpGs and facilitate the development of a more powerful epigenetic biomarker of aging. Using a innovative two-step process, we develop a new epigenetic biomarker of aging, DNAm PhenoAge, that strongly outperforms previous measures in regards to predictions for a variety of aging outcomes, including all-cause mortality, cancers, healthspan, physical functioning, and Alzheimer’s disease. While this biomarker was developed using data from whole blood, it correlates strongly with age in every tissue and cell tested. Based on an in-depth transcriptional analysis in sorted cells, we find that increased epigenetic, relative to chronological age, is associated increased activation of pro-inflammatory and interferon pathways, and decreased activation of transcriptional/translational machinery, DNA damage response, and mitochondrial signatures. Overall, this single epigenetic biomarker of aging is able to capture risks for an array of diverse outcomes across multiple tissues and cells, and provide insight into important pathways in aging.

## BACKGROUND

One of the major goals of geroscience research is to define ‘biomarkers of aging’[1, 2], which can be thought of as individual-level measures of aging that capture between-person differences in the timing of disease onset, functional decline, and death over the life course. While chronological age is arguably the strongest risk factor for aging-related death and disease, it is important to distinguish chronological time from biological aging. Individuals of the same chronological age may exhibit greatly different susceptibilities to age-related diseases and death, which is likely reflective of differences in their underlying biological aging processes. Such biomarkers of aging will be crucial to enable evaluation of interventions aimed at promoting healthier aging, by providing a measurable outcome, that unlike incidence of death and/or disease, does not require extremely long follow-up observation.

One potential biomarker that has gained significant interest in recent years is DNA methylation (DNAm). Chronological time has been shown to elicit predictable hypo-and hyper-methylation changes at many regions across the genome [3-7], and as a result, the first generation of DNAm based biomarkers of aging were developed to predict chronological age [8-13]. The blood-based algorithm by Hannum[10] and the multi-tissue algorithm by Horvath[14] produce age estimates (DNAm age) that correlate with chronological age well above r=0.90 for full age range samples. Nevertheless, while the current epigenetic age estimators exhibit statistically significant associations with many age-related diseases and conditions [15-26], the effect sizes are typically small to moderate. One explanation is that by using chronological age as the reference, by definition, may exclude CpGs whose methylation patterns don’t display strong time-dependent changes, but instead signal the departure of biological age from chronological age. Thus, it is important to not only capture CpGs that display changes with chronological time, but also those that account for differences in risk and physiological status among individuals of the same chronological age.

Previous work by us and others have shown that “phenotypic aging measures”, derived from clinical biomarkers[27-31], strongly predict differences in the risk of all-cause specific mortality, cause-specific mortality, physical functioning, cognitive performance measures, and facial aging among same-aged individuals. What’s more, in representative population data, some of these measures have been shown to be better indicators of remaining life expectancy than chronological age[27], suggesting that they may be approximating individual-level differences in biological aging rates. As a result, we hypothesize that a more powerful epigenetic biomarker of aging could be generated by replacing prediction of chronological age with prediction of a surrogate measure of “phenotypic aging” that, in and of itself, differentiates morbidity and mortality risk among same-age individuals.

## RESULTS

### Overview of the statistical model and analysis

Our development of the new epigenetic biomarker of aging proceeded along three main steps (Fig. 1). In step 1, a novel measure of ‘phenotypic age’ was developed using clinical data from the third National Health and Nutrition Examination Survey (NHANES). In step 2, DNAm from whole blood was used to predict phenotypic age, such that:

**Fig. 1.**
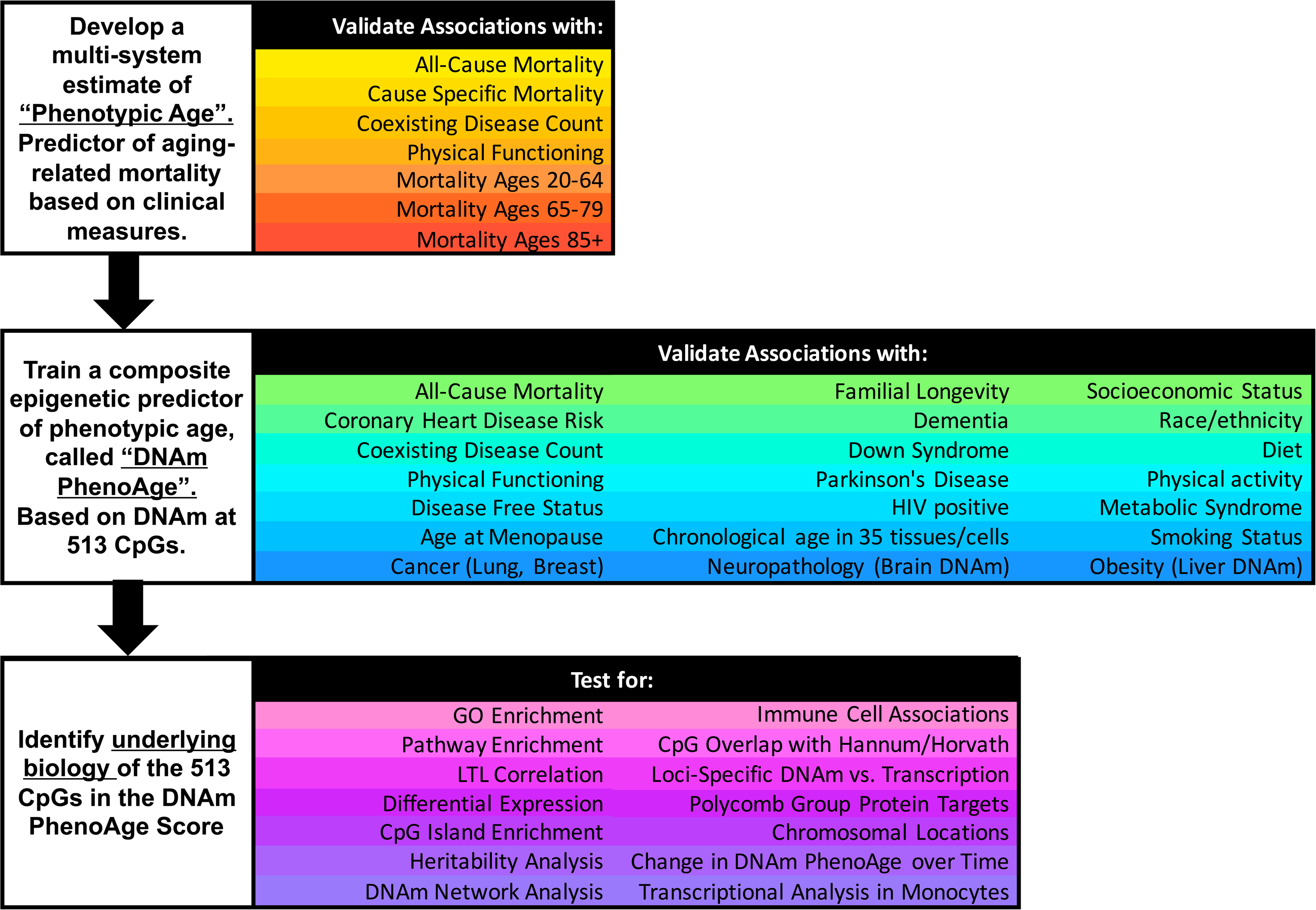
Roadmap for developing DNAm PhenoAge. The roadmap depicts our analytical procedures. In step 1, we developed an estimate of ‘Phenotypic Age’ based on clinical measure. Phenotypic age was developed using the NHANES III as training data, in which we employed a proportional hazard penalized regression model to narrow 42 biomarkers to 9 biomarkers and chronological age. This measure was then validated in NHANES IV and shown to be a strong predictor of both morbidity and mortality risk. In step 2, we developed an epigenetic biomarker of phenotypic age, which we call DNAm PhenoAge, by regressing phenotypic age (from step 1) on blood DNA methylation data, using the InCHIANTI data. This produced an estimate of DNAm PhenoAge based on 513 CpGs. We then validated our new epigenetic biomarker of aging, DNAm PhenoAge, using multiple cohorts, aging-related outcomes, and tissues/cells. In step 3, we examined the underlying biology of the 513 CpGs and the composite DNAm PhenoAge measure, using a variety of complementary data (gene expression, blood cell counts) and various genome annotation tools including chromatin state analysis and gene ontology enrichment.

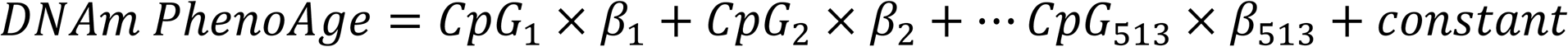

Predicted estimates from this model represent a person’s epigenetic age, which we refer to as ‘DNAm PhenoAge’. Using multiple independent datasets, we then tested whether DNAm PhenoAge was associated with a number of aging-related outcomes. We also tested whether it differed as a function of social, behavioral, and demographic characteristics, as whether it was applicable to tissues other than whole blood. Finally, in step 3, we examine the underlying biology of the 513 CpGs in the DNAm PhenoAge measure by examining differential expression, GO and pathway enrichment, chromosomal locations, and heritability.

### Estimating phenotypic age from clinical biomarkers

For step 1, NHANES III was used to generate a measure of phenotypic age. NHANES III is a nationally-representative sample, with over twenty-three years of mortality follow-up, from which our analytical sample included 9,926 adults with complete biomarker data. A Cox penalized regression model—where the hazard of mortality was regressed on forty-two clinical markers and chronological age—was used to select variables for inclusion in our phenotypic age score. The forty-two biomarkers considered represent those that were available in both NHANES III and IV. Based on 10-fold cross-validation, ten variables (including chronological age) were selected for the phenotypic age predictor (**Additional file 1: Table S1**). These nine biomarkers and chronological age were then combined in a phenotypic age estimate (in units of years) as detailed in Methods.

Validation data for phenotypic age came from NHANES IV, and included up to 17 years of mortality follow-up for n=6,209 national representative US adults. In this population, phenotypic age is correlated with chronological age at r=0.94. Results from all-cause and cause-specific (competing risk) mortality predictions, adjusting for chronological age (Table 1), show that a one year increase in phenotypic age is associated with a 9% increase in the risk of all-cause mortality (HR=1.09, p=3.8E-49), a 9% increase in the risk of mortality from aging-related diseases (HR=1.09, p=4.5E-34), a 10% increase in the risk of CVD mortality (HR=1.10, p=5.1E-17), a 7% increase in the risk of cancer mortality (HR=1.07, p=7.9E-10), a 20% increase in the risk of diabetes mortality (HR=1.20, p=1.9E-11), and a 9% increase in the risk of lung disease mortality (HR=1.09, p=6.3E-4). Further, phenotypic age is highly associated with comorbidity count (p=3.9E-21) and physical functioning measures (p=2.1E-10, Additional file 1: Fig. S1).

**Table 1:**
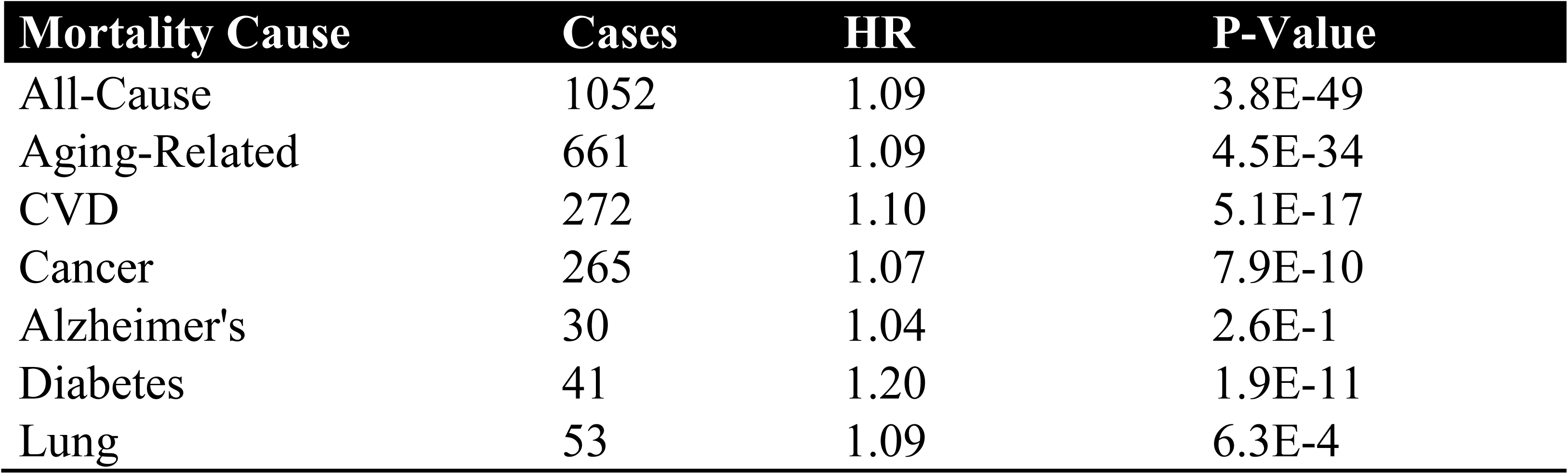
Mortality Validations for Phenotypic Age

| Mortality Cause | Cases | HR | P-Value |
| --- | --- | --- | --- |
| All-Cause | 1052 | 1.09 | 3.8E-49 |
| Aging-Related | 661 | 1.09 | 4.5E-34 |
| CVD | 272 | 1.10 | 5.1E-17 |
| Cancer | 265 | 1.07 | 7.9E-10 |
| Alzheimer's | 30 | 1.04 | 2.6E-1 |
| Diabetes | 41 | 1.20 | 1.9E-11 |
| Lung | 53 | 1.09 | 6.3E-4 |

### An epigenetic biomarker of aging (DNAm PhenoAge)

For step 2 (Fig. 1), data from n=456 participants at two time-points in the Invecchiare in Chianti (InCHIANTI) study was used to relate blood DNAm levels to phenotypic age. InCHIANTI was used as training data for the new epigenetic biomarker because the study assessed all clinical measures needed to estimate phenotypic age, contained data on DNAm, and had a large age range population (21-100 years). A total of 20,169 CpGs were considered when generating the new DNAm measure. They represented those CpGs available on all three chips (27k, 450k, EPIC), so as to facilitate usability across platforms. Elastic net regression, with 10-fold cross-validation, produced a model in which phenotypic age is predicted by DNAm levels at 513 of the 20,169 CpGs. The linear combination of the weighted 513 CpGs yields a DNAm based estimator of phenotypic age that we refer to as ‘DNAm PhenoAge’ (mean=58.9, s.d.=18.2, range=9.1-106.1), in contrast to the previously published Hannum and Horvath ‘DNAm Age’ measures.

While our new clock was trained on cross-sectional data in InCHIANTI, we capitalized on the repeated time-points to test whether changes in DNAm PhenoAge are related to changes in phenotypic age. As expected, between 1998 and 2007, mean change in DNAm PhenoAge was 8.51 years, whereas mean change in phenotypic age was 8.88 years. Moreover, participants’ phenotypic age (adjusting for chronological age) at the two time-points was correlated at r=0.50, whereas participants’ DNAm PhenoAge (adjusting for chronological age) at the two time-points was correlated at r=0.68 (**Additional file 1: Fig. S2**). Finally, we find that the change in phenotypic age between 1998 and 2007 is highly correlated with the change in DNAm PhenoAge between these two time-points (r=0.74, p=3.2E-80, **Additional file 1: Fig. S2**).

### DNAm PhenoAge strongly relates to all-cause mortality

In step 3 (Fig. 1), the epigenetic biomarker, DNAm PhenoAge, was calculated in five independent large-scale samples—two samples from Women’s Health Initiative (WHI) (n=2,016; and n=2,191), the Framingham Heart Study (FHS) (n=2,553), the Normative Aging Study (NAS) (n=657), and the Jackson Heart Study (JHS) (n=1,747). The first four studies used the Illumina 450K array while the JHS employed the latest Illumina EPIC array platform. In these studies, DNAm PhenoAge correlated with chronological age at r=0.66 in WHI (Sample 1), r=0.69 in WHI (Sample 2), r=0.78 in FHS, r=0.62 in the NAS, and r=0.89 in JHS. The five validation samples were then used to assess the effects of DNAm PhenoAge on mortality in comparison to the Horvath and Hannum DNAm Age measures. DNAm PhenoAge was significantly associated with subsequent mortality risk in all studies (independent of chronological age), such that, a one year increase in DNAm PhenoAge is associated with a 4.5% increase in the risk of all-cause mortality (Meta(FE)=1.045, Meta p=7.9E-47, Fig. 2). To better conceptualize what this increase represents, we compared the predicted life expectancy and mortality risk for person’s representing the top 5% (fastest agers), the average, and the bottom 5% (slowest agers). Results suggest that those in the top 5% of fastest agers have a mortality hazard of death that is about 1.62 times that of the average person, i.e. your hazard of death is 62% higher than that of an average person. Further, contrasting the 5% fastest agers with the 5% slowest agers, we find that the hazard of death of the fastest agers is 2.58 times higher than that of the bottom 5% slowest agers (HR=1.045^11.0^/1.045^−10.5^). Additionally, both observed and predicted Kaplan-Meier survival estimates showed that faster agers had much lower life expectancy and survival rates compared to average and/or slow agers (Fig. 2).

**Fig. 2.**
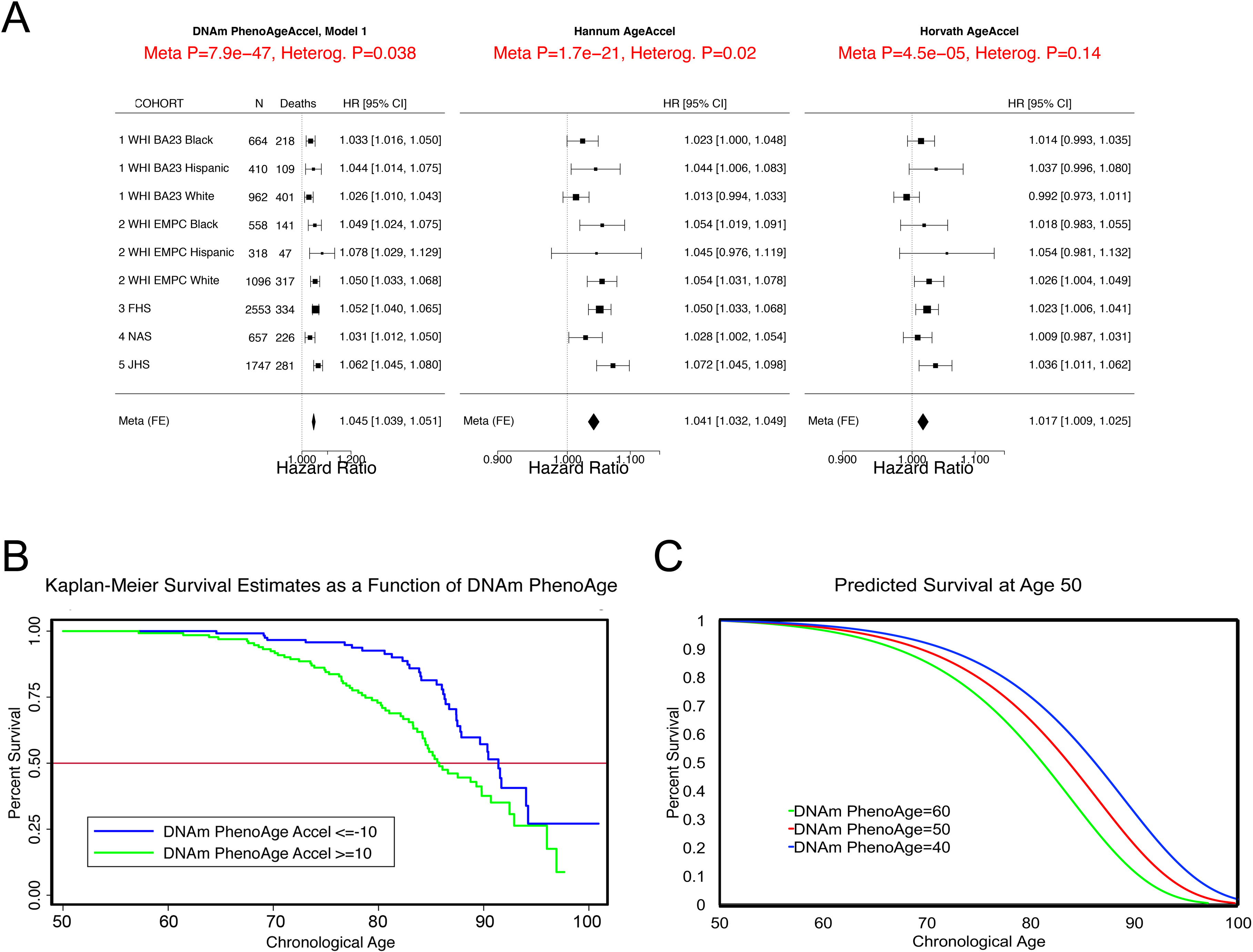
Mortality Prediction by DNAm PhenoAge. A: Using five samples from large epidemiological cohorts—two samples from the Women’s health Initiative, the Framingham Heart Study, the Normative Aging Study, and the Jackson Heart Study—we tested whether DNAm PhenoAge was predictive of all-cause mortality. The Fig. displays a forest plot for fixed-effect meta-analysis, based on Cox proportional hazard models, and adjusting for chronological age. Results suggest that DNAm PhenoAge is predictive of mortality in all samples, and that overall, a one year increase in DNAm PhenoAge is associated with a 4.5% increase in the risk of death (p=9.9E-47). This is contrasted against the first generation of epigenetic biomarkers of aging by Hannum and Horvath, which exhibit less significant associations with lifespan (p=1.7E-21 and p=4.5E-5, respectively). B & C: Using the WHI sample 1, we plotted Kaplan-Meier survival estimates using actual data from the fastest versus the slowest agers (panel B). We also applied the equation from the proportional hazard model to predict remaining life expectancy and plotted predicted survival assuming a chronological age of 50 and a DNAm PhenoAge of either 40 (slow ager), 50 (average ager), or 60 (fast ager) (panel C). Median life expectancy at age 50 was predicted to be approximately 81 years for the fastest agers, 83.5 years for average agers, and 86 years for the slowest agers.

As shown in Fig. 2, the DNAm age based measures from Hannum and Horvath also related to all-cause mortality, consistent with what has been reported previously [16, 24, 32-34]. To directly compare the three epigenetic measures, we contrasted their accuracy in predicting 10-year and 20-year mortality risk, using receiver operating characteristics curves. DNAm PhenoAge (adjusted for age) predicts both 10-year mortality and 20-year mortality significantly better than the other two measures (Additional file 1: Table S2). Finally, when examining a model that includes all three measures (Additional file 1: Table S3), we find that only DNAm PhenoAge is positively associated with mortality (HR=1.04, p=1.33E-8), whereas Horvath DNAm Age is now negatively associated (HR=0.98, p=2.72E-2), and Hannum DNAm Age has no association (HR=1.01, p=4.66E-1).

### DNAm PhenoAge strongly relates to aging-related morbidity

Given that aging is believed to also influence diseases incidence/prevalence, we examined whether DNAm PhenoAge relate to diverse ag-related morbidity outcomes. We observe strong associations between DNAm PhenoAge and a variety of other aging outcomes using the same five validation samples (Table 2). For instance, independent of chronological age, higher DNAm PhenoAge is associated with an increase in a person’s number of coexisting morbidities (β=0.008 to 0.031; Meta P-value=1.95E-20), a decrease in likelihood of being disease-free (β=−0.002 to −0.039; Meta P-value=2.10E-10), an increase in physical functioning problems (β=-0.016 to −0.473; Meta P-value=2.05E-13), an increase in the risk of CHD risk (β=0.016 to 0.073; Meta P-value=3.35E-11).

**Table 2:**
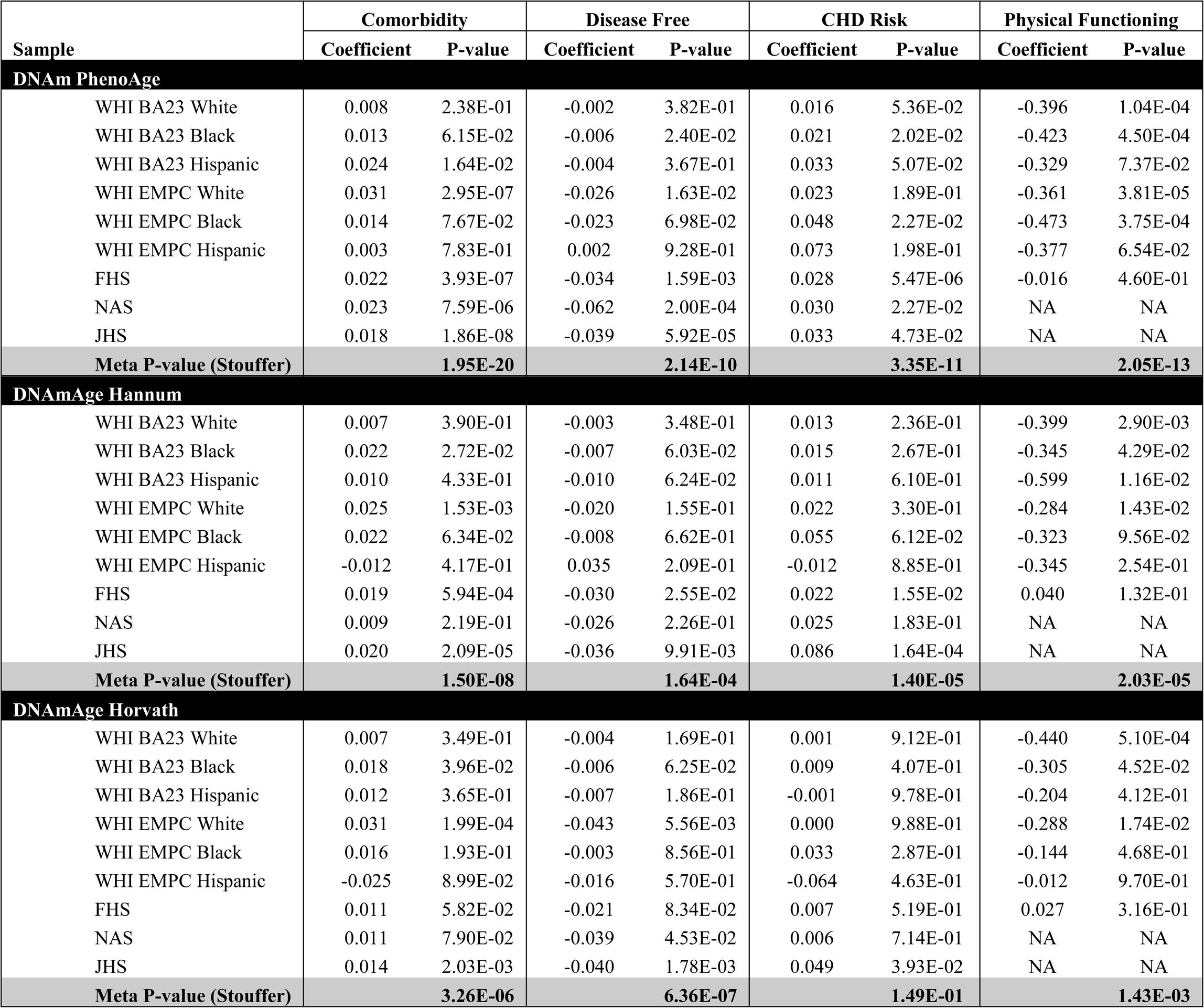
Morbidity Validation for DNAm PhenoAge

### DNAm PhenoAge and smoking

Cigarette exposure has been shown to have an epigenetic fingerprint[35-37], which has been reflected in previous DNAm risk predictors[38]. Similarly, we find that DNAm PhenoAge significantly differs between never (n=1,097), current (n=209), and former smokers (n=710) (p=0.0033) (Additional file 1, Fig. S3A); however, conversely, we do not find a robust association between pack-years and DNAm PhenoAge (Additional file 1, Fig. S3B-D). Given the association between DNAm PhenoAge and smoking, we re-evaluated the morbidity and mortality associations (fully-adjusted) in our four samples, stratifying by smoking status (Additional file 1: Fig. S4 and Table S4). We find that DNAm PhenoAge is associated with mortality among both smokers (adjusted for pack-years) (Meta(FE)=1.050, Meta p=7.9E-31), and non-smokers (Meta(FE)=1.033, Meta p=1.2E-10). DNAm PhenoAge relates to the number of coexisting morbidities, physical functioning status, disease free status, and CHD for both smokers and non-smokers (Additional file 1: Table S4). Finally, in previous work we showed that Horvath DNAm age of blood predicts lung cancer risk in the first WHI sample[21]. Using the same data, we find that a one year increase in DNAm PhenoAge (adjusting for chronological age, race/ethnicity, pack-years, and smoking status) is associated with a 5% increase in the risk lung cancer incidence and/or mortality (HR=1.05, p=0.031). Further, when restricting the model to current smokers only, we find that the effect of DNAm PhenoAge on future lung cancer incidence and/or mortality is even stronger (HR=1.10, p=0.014).

In evaluating the relationship between DNAm PhenoAge and additional characteristics we observe significant differences between racial/ethnic groups (p=5.1E-5), with non-Hispanic blacks having the highest DNAm PhenoAge on average, and non-Hispanic whites having the lowest (Additional file 1: Fig. S5). We also find evidence of social gradients in DNAm PhenoAge, such that those with higher education (p=6E-9) and higher income (p=9E-5) appear younger. DNAm PhenoAge relates to exercise and dietary habits, such that increased exercise (p=7E-5) and markers of fruit/vegetable consumption (such as carotenoids, p=2E-27) are associated with lower DNAm PhenoAge (Additional file 1: Fig. S6A & Additional file 1: Fig. S6B). Cross sectional studies in the WHI also revealed that DNAmPhenoAge acceleration is positively correlated with C-reactive protein (r=0.18, p=5E-22, Additional file 1: Fig. S6B), insulin (r=0.15, p=2E-20), glucose (r=0.10, p=2E-10), triglycerides (r=0.09, p=5E-9), waist to hip ratio (r=0.15, p=5E-22) but it is negatively correlated with the “good” cholesterol HDL (r=-0.09, p=7E-9).

### DNAm PhenoAge in other tissues

One advantage of developing biological aging estimates based on molecular markers (like DNAm), rather than clinical risk measures (e.g. those in the phenotypic age variable), is that they may lend themselves to measuring tissue/cell specific aging. Although DNAm PhenoAge was developed using samples from whole blood, our empirical results show that it strongly correlates with chronological age in a host of different tissues and cell types (Fig. 3). For instance, when examining all tissues concurrently, the correlation between DNAm PhenoAge and chronological age was 0.71. Age correlations in brain tissue ranged from 0.54 to 0.92, while correlations were also found in breast (r=0.47), buccal cells (r=0.88), dermal fibroblasts (r=0.87), epidermis (r=0.84), colon (r=0.88), heart (r=0.66), kidney (r=0.64), liver (r=0.80), lung (r=055), and saliva (r=0.81).

**Fig. 3.**
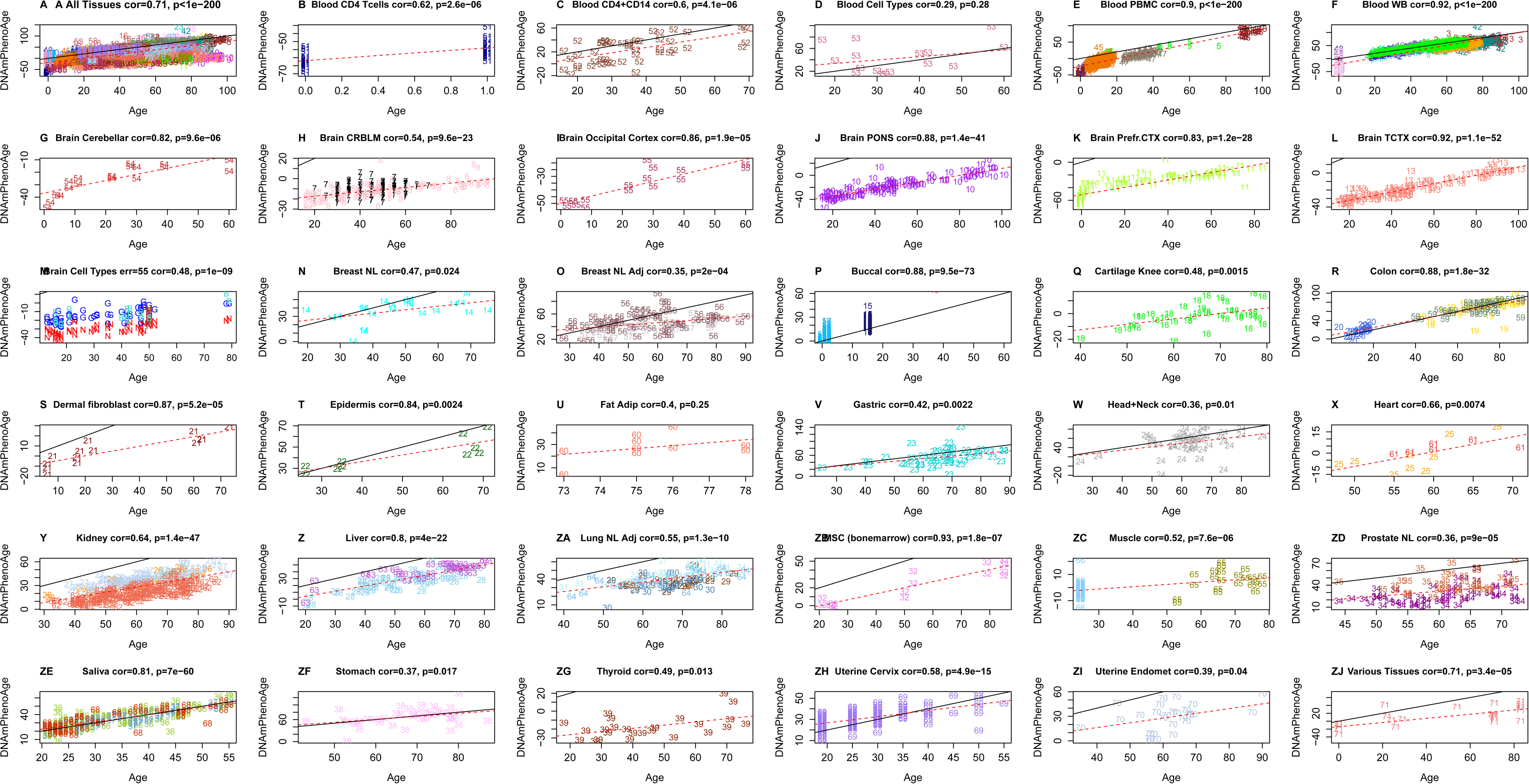
Chronological age versus DNAm PhenoAge in a variety of tissues and cells. Although DNAm PhenoAge was developed using methylation data from whole blood, it also tracks chronological age in a wide variety of tissues and cells. A) The correlation across all tissues/cells we examined is r=0.71. B-ZJ) report results in different sources of DNA as indicated in panel headings. The numbers correspond to the data sets from (Horvath 2013). Overall, correlations range from r=0.35 (breast, panel O) to r=0.92 (temporal cortex in brain, panel L).

### Alzheimer’s disease and brain samples

Based on the accuracy of the age prediction in other tissues/cells, we examined whether aging in a given tissue was associated with tissue-associated outcomes. For instance, using data from approximately 700 post-mortem samples from the Religious Order Study (ROS) and the Memory and Aging Project (MAP)[39, 40] we tested the association between pathologically diagnosed Alzheimer’s disease and DNAm PhenoAge in dorsolateral prefrontal cortex (DLPFX). Results suggest (Fig. 4) that those who are diagnosed with Alzheimer’s disease (AD), based on postmortem autopsy, have DLPFX that appear more than one year older than same aged individuals who are not diagnosed with AD postmortem (p=4.6E-4). Further, age adjusted DNAm PhenoAge was found to be positively associated with neuropathological hallmarks of Alzheimer’s disease, such as amyloid load (r=0.094, p=0.012), neuritic plaques (r=0.11, p=0.0032), and neurofibrillary tangles (r=0.10, p=0.0073).

**Fig. 4.**
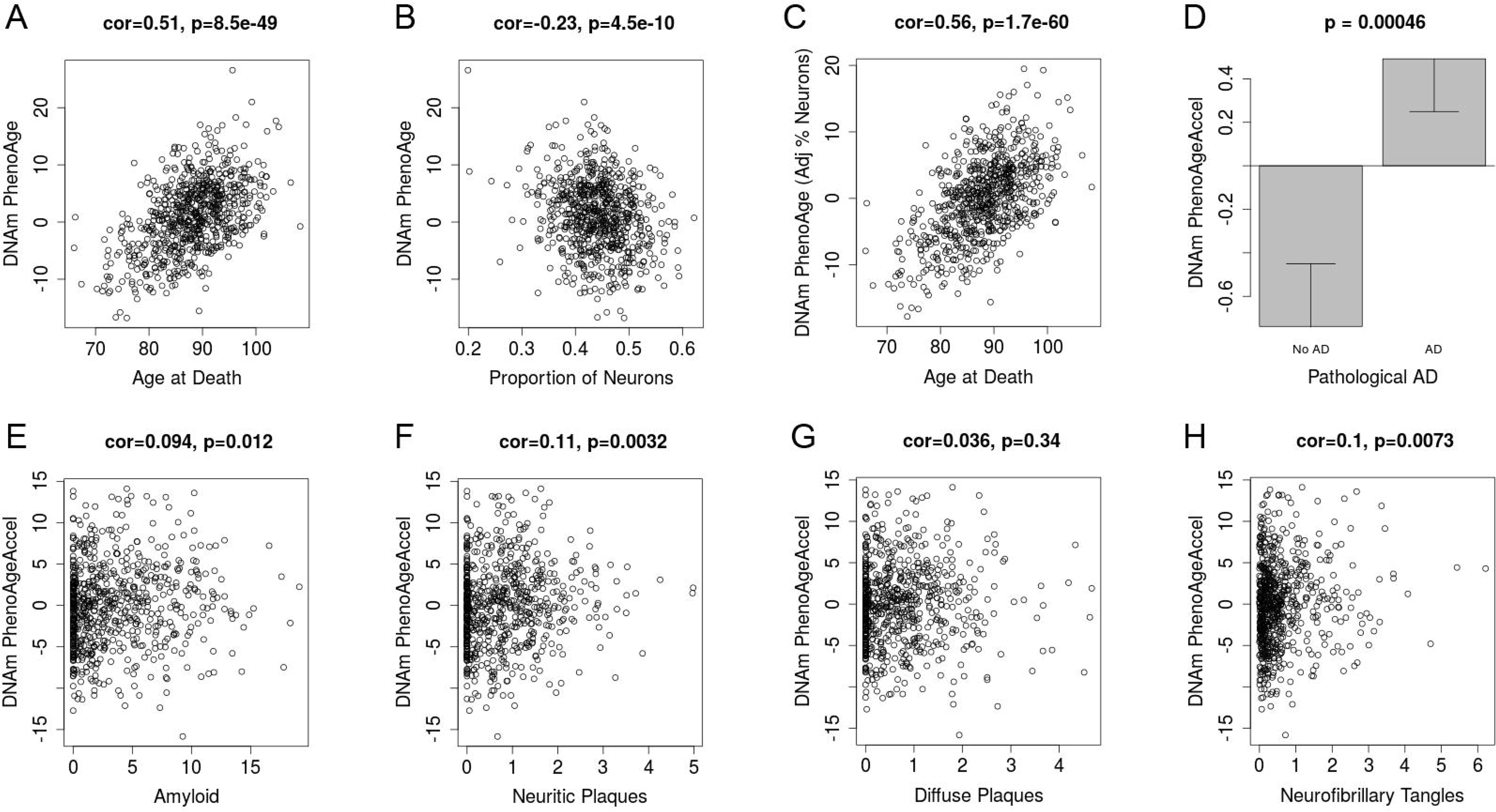
DNAm PhenoAge measured in dorsolateral prefrontal cortex relates to Alzheimer’s disease and related neuropathologies. Using postmortem data from the Religious Order Study (ROS) and the Memory and Aging Project (MAP), we find a moderate/high correlation between chronological age and DNAm PhenoAge (panl A), that is further increased after adjusting for the estimated proportion on neurons in each sample (panel C). We also find that DNAm PhenoAge is significantly higher (p=0.00046) among those with Alzheimer’s disease versus controls (panel D), and that it positively correlates with amyloid load (p=0.012, panel E), neuritic plaques (p=0.0032, panel F), diffuse plaques (p=0.036, panel G), and neurofibrillary tangles (p=0.0073, panel H).

### Comparison with other DNAm biomarkers of aging

Several additional DNAm biomarkers have been described in the literature CpGs [12, 13]. A direct comparison of 6 DNAm biomarkers (including DNAm PhenoAge) reveals that DNAm PhenoAge stands out in terms of its predictive accuracy for lifespan, its relationship with smoking status, its relationship with leukocyte telomere length, naïve CD8+ T cells and CD4+ T cells (Additional file 1: Table S5).

### DNAm PhenoAge and Immunosenescence

To test the hypothesis that DNAm PhenoAge captures aspects of the age-related decline of the immune system, we correlated DNAm PhenoAge with estimated blood cell count (Additional file 1, Fig. S7). After adjusting for age, we find that DNAm PhenoAgeAccel is negatively correlated with naïve CD8+ T cells (r=-0.35, p=9.2E-65), naïve CD4+ T cells (r=-0.29, p=4.2E-42), CD4+ helper T cells (r=−0.34, p=3.6E-58), and B cells (r=−0.18, p=8.4E-17). Further, DNAm PhenoAgeAccel is positively correlated with the proportion of granulocytes (r=0.32, p=2.3E-51), exhausted CD8+ (defined as CD28-CD45RA-) T cells (r=0.20, p=1.9E-20), and plasma blast cells (r=0.26, p=6.7E-34). These results are consistent with age related changes in blood cells [41] and suggest that DNAm PhenoAge may capture aspects of immunosenescence in blood. However, three lines of evidence suggest that DNAm PhenoAge is not simply a measure of immunosenescence. First, another measure of immunosenescence, leukocyte telomere length, is only weakly correlated with DNAm PhenoAgeAccel (r=-0.13 p=0.00019 in the WHI and r=-0.087, P=7.6E-3 in Framingham Heart study, Additional file 1, Fig. S8). Second, the strong association between DNAm PhenoAge and mortality does not simply reflect changes in blood cell composition, as can be seen from the fact that in Additional file 1, Fig. S9 the robust association remains even after adjusting for estimates of seven blood cell count measures (Meta(FE)=1.036, Meta p=5.6E-21).

### DNA sequence characteristics of the 513 CpGs in DNAm PhenoAge

Of the 513 CpGs in DNAm PhenoAge, we find that, 41 CpGs were also in the Horvath DNAm Age measure (Additional file 2: Table S6). This represents a 4.88-fold increase over what would be expected by chance (p=8.97E-15). Of the 41 overlapping CpGs, the average absolute value for their age correlations was r=0.40, and 31 had age correlations with absolute values in the top 20% of what is found for among the 513 CpGs in the DNAm PhenoAge score. We also observed 6 CpGs that overlapped between the Hannum DNAm Age score and the DNAm PhenoAge score—five of which were also found in the Horvath DNAm Age measure. All six CpGs had extremely high age correlations (half positive, half negative), with absolute values between r=0.49 and r=0.76. The five CpGs that are found in all three epigenetic aging measures were: cg05442902 (*P2RXL1*), cg06493994 (*SCGN*), cg09809672 (*EDARADD*), cg19722847 (*IPO8*), and cg22736354 (*NHLRC1*).

Finally, we conducted a functional enrichment analysis of the chromosomal locations of the 513 CpGs, we found that 149 CpGs whose age correlation exceeded 0.2 tended to be located in CpG islands (p=0.0045, Additional file 1: Fig. S10) and were significantly enriched with polycomb group protein targets (p=8.7E-5, Additional file 1: Fig. S10), which echoes results of epigenome wide studies of aging effects [4, 42, 43].

### Transcriptional and genetic studies of DNAm PhenoAge

Using the genome-wide data from FHS and WHI, we estimated the heritability of DNAm PhenoAge. The heritability estimated by the SOLAR polygenic model for those of European ancestry in the FHS was *h*^2^=0.33, while the heritability estimated for those of European ancestry in WHI, using GCTA-GREML analysis[44, 45] was *h*^2^=0.51.

Using the monocyte data mentioned above, as well as PBMC expression data on 2,188 persons from the FHS, we conducted a more thorough transcriptional analysis to identify differential expression associated with DNAm PhenoAgeAccel (Additional file 3: Table S7). Overall, we find that genes show similar associations to chronological age and DNAm PhenoAgeAccel. DNAm PhenoAgeAccel represents aging differences among same-aged individuals and is adjusted so as to exhibit a correlation of r=0.0 with chronological age. Thus, this observation can be taken to suggest that genes whose transcription increases with age are upregulated among epigenetically older compared to epigenetically younger persons of the same chronological age (Additional file 1: Fig. S11)—same applies for genes that show decreases with chronological age being downregulated in epigenetically older versus younger persons of the same age.

Using the transcriptional data from monocytes described above (adjusting for array, sex, race/ethnicity, age, and imputed cell counts), we tested for GO enrichment among genes that positively associated with DNAm PhenoAge and those that negatively associated with DNAm PhenoAge (Additional file 4: Table S8). Among those with positive aging associations (overexpression among epigenetically older individuals), we observed enrichment for a number of pro-inflammatory signaling pathways. These pathways included, but were not limited to: multiple toll-like receptor signaling pathways (7,9,3,2), regulation of inflammatory response, JAK-STAT cascade, response to lipopolysaccharide, tumor necrosis factor-mediated signaling pathway, and positive regulation of NF-kappaB transcription factor activity. Additionally, positively associated genes were also enriched for a number anti-viral response pathways—type I interferon signaling, defense response to virus, interferon-gamma-mediated signaling pathway, cellular response to interferon-alpha, etc. Finally, other interesting GO terms enriched among positively associated genes included: response to nutrient, JAK-STAT cascade involved in growth hormone signaling pathway, multicellular organism growth, and regulation of DNA methylation.

When testing for enrichment among genes that were negatively associated with DNAm PhenoAgeAccel (decreased expression among epigenetically older persons) we observed that many were implicated in processes involving transcriptional and translational machinery, as well as damage recognition and repair. These included: translational initiation; regulation of translational initiation; ribosomal large subunit assembly; ribosomal small subunit assembly; translational elongation; transcription initiation from RNA polymerase I promoter; transcription-coupled nucleotide-excision repair; nucleotide-excision repair, DNA incision, 5’-to lesion; nucleotide-excision repair, DNA damage recognition; DNA damage response, detection of DNA damage; and regulation of DNA damage checkpoint.

## DISCUSSION

Using a novel two-step method, we were successful in developing a DNAm based biomarker of aging that is highly predictive of nearly every morbidity and mortality outcome we tested. Training an epigenetic predictor of phenotypic age instead of chronological age led to substantial improvement in mortality/healthspan predictions over the first generation of DNAm based biomarkers of chronological age from Hannum[10], Horvath[14] and other published DNAm biomarkers. In doing so, this is the first study to conclusively demonstrate that DNAm biomarkers of aging are highly predictive of CVD and coronary heart disease. DNAm PhenoAge also tracks chronological age and relates to disease risk in samples other than whole blood. Finally, we find that an individual’s DNAm PhenoAge, relative to his/her chronological age, is moderately heritable and is associated with activation of pro-inflammatory, interferon, DNAm damage repair, transcriptional/translational signaling, and various markers of immunosenescenc: a decline of naïve T cells and shortened leukocyte telomere length.

The ability of our measure to predict multifactorial aging conditions is consistent with the fundamental underpinnings of Geroscience research [1, 46], which posits that aging mechanisms give rise to multiple pathologies and thus, differences in the rate of aging will have implications for a wide array of diseases and conditions. Further, these results answer a fundamental biological question of whether differences in multi-system dysregulation (estimated using clinical phenotypic age measures), healthspan, and lifespan are reflected at the epigenetic level, in the form of differential DNAm at specific CpG sites.

The improvement over previous epigenetic biomarkers, likely comes down to the types of CpGs selected for the various measures. Only 41 of the 513 CpGs in DNAm PhenoAge were shared with the Horvath clock, while only five CpGs were shared between all three clocks (DNAm PhenoAge, Horvath, and Hannum). In general, these CpGs did not tend to be drivers of the DNAm PhenoAge score, and instead represented those with large age correlations, but lower weights. This may explain the improvements of DNAm PhenoAge over previous epigenetic biomarkers of aging. While the previous DNAm age estimators selected CpGs to optimize prediction of chronological age, the CpGs in DNAm PhenoAge were optimized to predict a multi-system proxy of physiological dysregulation (phenotypic age). In doing so, we were able to not only capture CpGs that exhibited changes in DNAm with age, but also those that captured variations in risk of death and disease among same aged individuals. In general, the CpGs with the highest weights in the new clock did not correlate with chronological age (Additional file 1: Fig. S12), but instead were related to the difference between phenotypic and chronological age—i.e. divergence in the rate of aging. Interestingly, the CpGs that contributed the most to the DNAm PhenoAge score, tended to have low age correlations.

While DNAm PhenoAge greatly outperformed all previous DNAm biomarkers of aging (Additional file 1: Table S5), the utility of DNAm PhenoAge for estimating risk does not imply that it should replace clinical biomarkers when it comes to informing medical and health-related decisions. In fact, but perhaps not surprisingly, the phenotypic age measure used to select CpGs is a better predictor of morbidity and mortality outcomes than DNAm PhenoAge. While the addition of error in performing a two-step process, rather than training a DNAm predictor directly on mortality may contribute, we don’t believe this accounts for the difference in predictive performance. In fact, a recent DNAm measure by Zhang et al.[38] was trained to directly predict mortality risk, yet it appears to be a weaker predictor than both our DNAm PhenoAge measure and our clinical phenotypic age measure (Additional file 1: Table S9). The first generation of DNAm age estimators only exhibit weak associations with clinical measures of physiological dysregulation [47, 48]. Physiological dysregulation, which is more closely related to our clinical age measure “phenotypic age” than to chronological age, is not only the result of exogenous/endogenous stress factors (such as obesity, infections) but also a result of age related molecular alterations, one example of which are modifications to the epigenome. Over time, dysregulation within organ systems leads to pathogenesis of disease (age-related molecular changes → physiological dysregulation → morbidity → mortality)[49]. However, stochasticity and variability exist at each of these transitions. Therefore, measures of physiological dysregulation, will be better predictors of transition to the next stage in the aging trajectory (i.e. morbidity and mortality) than will measures of age related molecular alterations, like DNAm PhenoAge. Similarly, quantification of disease pathogenesis (cancer stage, Alzheimer’s stage) is likely a better predictor of mortality risk than clinical phenotypic aging measures. As a result, clinical phenotypic aging measures may be preferable to epigenetic measures when the goal is risk prediction, and all samples come from blood.

That being said, when the aim is to study the mechanisms of the aging process, DNAm measures have advantages over clinical measures. First, they may better capture “pre-clinical aging” and thus may be more suited for differentiating aging in children, young adults, or extremely healthy individuals, for whom measures like CRP, albumin, creatinine, glucose, etc. are still fairly homogenous. Second, as demonstrated, these molecular measures can capture cell and/or tissue specific aging rates and therefore may also lend themselves to in vitro studies of aging, studies for which blood is not available, studies using postmortem samples, and/or studies comparing aging rates between tissues/cells. While the fundamental drivers of aging are believed to be shared across cells/tissues, that is not to say that all the cells and tissues within an individual will age at the same rate. In fact, it is more likely that individuals will vary in their patterning of aging rates across tissues, and that this will have implications for death and disease risk. Relatedly, it is not known how predictions based on DNAm PhenoAge measures from non-blood samples will compare to phenotypic age predictions. It may be the case that various outcomes will be more tightly related to aging in specific cells/tissues, rather than blood. Finally, examination of DNAm based aging rates facilitates the direct study of the proposed mechanisms of aging, of which “epigenetic alterations” is one of the seven hypothesized “pillars of aging” [1].

While more work needs to be done to model the biology linking DNAm PhenoAge and aging outcomes, we began to explore this using differential expression, functional enrichment, heritability estimates, and network analysis. Overall, we found that CpGs that had larger increases with aging tended to be located in CpG islands and enriched with polycomb group protein targets, consistent with what has been reported in previous epigenome wide studies of aging effects [4, 6, 7, 42, 43]. While typically DNAm of CpG islands and/or polycomb recruitment is linked to transcriptional silencing [50], for the most part, we did not observe associations between DNAm and expression for co-locating CpG-gene pairs—this was also true when only considering CpGs located in islands. These findings may suggest that the genes annotated to the CpGs in our score are not part of the link between changes in DNAm and aging. Nevertheless, we also recognize that these null results could stem from the fact that 1) associations were only tested in monocytes, 2) DNAm and expression represents what is present globally for each sample, rather than on a cell-by-cell bases, and 3) stronger associations between DNAm and gene expression levels may only exist early in life.

Nevertheless, we do identify potentially promising transcriptional pathways when considering DNAm PhenoAge as a whole. For instance, we observe that higher DNAm PhenoAge is associated with increases in the activation of proinflammatory pathways, such as NF-kappaB; increased interferon (IFN) signaling; decreases in ribosomal–related and translational machinery pathways; and decreases in damage recognition and repair pathways. These findings are consistent with previous work describing aging associated changes, comprising increases in dysregulated inflammatory activation, increased DNA damage, and loss of translational fidelity. For instance, there exists a large body of literature highlighting the importance of an increased low-grade pro-inflammatory status as a driver of the aging process, termed inflamm-aging [41, 51, 52]. IFN signaling pathways have been shown to be markers of DNA damage and mediators of cellular senescence[53]. Additionally, it has been shown that breakdown of the transcriptional and translational machinery may play a central role in the aging process[54, 55]. For instance, the ribosome is believed to be a key regulator of proteostasis, and in turn, aging[54, 56]. Relatedly, loss of integrity in DNA damage repair pathways is considered another hallmark of the aging process[57-59].

In general, many of these pathways will have implications for adaptation to exogenous and endogenous stressors. Factors related to stress resistance and response have repeatedly been shown to be drivers of differences in lifespan and aging[60-65]. This may partially account for our findings related to smoking. In general, it is not surprising that a biomarker of aging and mortality risk relates to smoking, given that life expectancies of smokers are on average ten years shorter than never smokers, and smoking history is associated with a drastic increase in the risk of a number of age-related conditions. However, perhaps more interestingly, we find that the effects of DNAm PhenoAge on mortality appear to be higher for smokers than non-smokers, which could suggest that DNAm PhenoAge represent differences in innate resilience/vulnerability to pro-aging stressors, such as cigarette smoke.

Interestingly, we observed moderately high heritability estimates for DNAm PhenoAge. For instance, we estimated that genetic differences accounted for one-third to one-half of the variance in DNAm PhenoAge, relative to chronological age. In moving forward, it will be useful to identify the genetic architecture underlying differences in epigenetic aging. Finally, we reported that individuals’ DNAm PhenoAges—relative to their chronological ages—remained fairly stable over a nine-year period. However, it is unclear whether it is attributable to genetic influences, or the fact that social and behavioral characteristics tend to also remain stable for most individuals.

If the goal is to utilize accurate quantifiable measures of the rate of aging, such as DNAm PhenoAge, to assess the efficacy of aging interventions, more work will be needed to evaluate the dynamics of DNAmPhenoAge following various treatments. For instance, it remains to be seen whether interventions can reverse DNAmPhenoAge in the short term. Along these lines, it will be essential to determine causality—does DNAm drive the aging process, or is it simply a surrogate marker of senescence? If the former is true, DNAm PhenoAge could provide insight into promising targets for therapies aimed at lifespan, and more importantly, healthspan extension.

## Conclusion

Overall, DNAm PhenoAge is an attractive composite biomarker that captures organismal age and the functional state of many organ systems and tissues, above and beyond what is explained by chronological time. Our validation studies in multiple large and independent cohorts demonstrate that DNAm PhenoAge is a highly robust predictor of both morbidity and mortality outcomes, and represents a promising biomarker of aging, which may prove to be beneficial to both basic science and translational research.

## METHODS

Using the NHANES training data, we applied a Cox penalized regression model—where the hazard of aging-related mortality (mortality from diseases of the heart, malignant neoplasms, chronic lower respiratory disease, cerebrovascular disease, Alzheimer’s disease, Diabetes mellitus, nephritis, nephrotic syndrome, and nephrosis) was regressed on forty-two clinical markers and chronological age to select variables for inclusion in our phenotypic age score. Ten-fold cross-validation was employed to select the parameter value, lambda, for the penalized regression. In order to develop a sparse phenotypic age estimator (the fewest biomarker variables needed to produce robust results) we selected a lambda of 0.0192, which represented a one standard deviation increase over the lambda with minimum mean-squared error during cross-validation (Additional file 1, Fig. S13). Of the forty-two biomarkers included in the penalized Cox regression model, this resulted in ten variables (including chronological age) that were selected for the phenotypic age predictor.

These nine biomarkers and chronological age were then included in a parametric proportional hazards model based on the Gompertz distribution. Based on this model, we estimated the 10-year (120 months) mortality risk of the j-the individual. Next, the mortality score was converted into units of years (Additional file 1). The resulting phenotypic age estimate was regressed on DNA methylation data using an elastic net regression analysis. The penalization parameter was chosen to minimize the cross validated mean square error rate (Additional file 1, Fig. S14), which resulted in 513 CpGs.

### Estimation of blood cell counts based on DNAm levels

We estimate blood cell counts using two different software tools. First, Houseman’s estimation method [66] was used to estimate the proportions of CD8+ T cells, CD4+ T, natural killer, B cells, and granulocytes (mainly neutrophils). Second, the Horvath method, implemented in the advanced analysis option of the epigenetic clock software [11, 18], was used to estimate the percentage of exhausted CD8+ T cells (defined as CD28-CD45RA-), the number (count) of naïve CD8+ T cells (defined as CD45RA+CCR7+) and plasmablasts. We and others have shown that the estimated blood cell counts have moderately high correlations with corresponding flow cytometric measures [66, 67].

Additional descriptions of methods and materials can be found in Additional file 1.

## Ethics approval

This study was reviewed by the UCLA institutional review board (IRB#13-000671, IRB#15-000697, IRB#16-001841, IRB#15-000682)

## Author contributions

ML and SH developed the DNAmPhenoAge estimator and wrote the article. SH, ML, LF conceived of the study. ML, SH, AL, AQ carried out the statistical analysis. The remaining authors contributed data and participated in the interpretation of the results.

## ACKNOWLEDGEMENTS

We would like to acknowledge The WHI Investigators listed below:

Program Office: (National Heart, Lung, and Blood Institute, Bethesda, Maryland) Jacques Rossouw, Shari Ludlam, Dale Burwen, Joan McGowan, Leslie Ford, and Nancy Geller.

Clinical Coordinating Center: (Fred Hutchinson Cancer Research Center, Seattle, WA) Garnet Anderson, Ross Prentice, Andrea LaCroix, and Charles Kooperberg.

Investigators and Academic Centers: (Brigham and Women’s Hospital, Harvard Medical School, Boston, MA) JoAnn E. Manson; (MedStar Health Research Institute/Howard University, Washington, DC) Barbara V. Howard; (Stanford Prevention Research Center, Stanford, CA).

Marcia L. Stefanick; (The Ohio State University, Columbus, OH) Rebecca Jackson; (University of Arizona, Tucson/Phoenix, AZ) Cynthia A. Thomson; (University at Buffalo, Buffalo, NY) Jean Wactawski-Wende; (University of Florida, Gainesville/Jacksonville, FL) Marian Limacher; (University of Iowa, Iowa City/Davenport, IA) Robert Wallace; (University of Pittsburgh, Pittsburgh, PA) Lewis Kuller; (Wake Forest University School of Medicine, Winston-Salem, NC) Sally Shumaker.

The Framingham Heart Study is funded by National Institutes of Health contract N01-HC-25195 and HHSN268201500001I. The laboratory work for this investigation was funded by the Division of Intramural Research, National Heart, Lung, and Blood Institute, National Institutes of Health. The analytical component of this project was funded by the Division of Intramural Research, National Heart, Lung, and Blood Institute, and the Center for Information Technology, National Institutes of Health, Bethesda, MD. The Framingham Heart Study is conducted and supported by the National Heart, Lung, and Blood Institute (NHLBI) in collaboration with Boston University. This manuscript was not prepared in collaboration with investigators of the Framingham Heart Study and does not necessarily reflect the opinions or views of the Framingham Heart Study, Boston University, or the NHLBI.

The United States Department of Veterans Affairs (VA) Normative Aging Study (NAS) is supported by the Cooperative Studies Program/ERIC and is a research component of the Massachusetts Veterans Epidemiology Research and Information Center (MAVERIC), Boston Massachusetts.

The MESA Epigenomics and Transcriptomics Studies were funded by R01HL101250, R01 DK103531-01, R01 DK103531, R01 AG054474, and R01 HL135009-01 to Wake Forest University Health Sciences.

We thank the Jackson Heart Study (JHS) participants and staff for their contributions to this work. The JHS is supported by contracts HHSN268201300046C, HHSN268201300047C, HHSN268201300048C, HHSN268201300049C, HHSN268201300050C from the National Heart, Lung, and Blood Institute and the National Institute on Minority Health and Health Disparities. Dr. Wilson is supported by U54GM115428 from the National Institute of General Medical Sciences.

## FUNDING

This study was supported by NIH/NIA U34AG051425-01 (Horvath) and NIH/NIA K99AG052604 (Levine). The WHI epigenetic studies were supported by NIH/NHLBI 60442456 BAA23 (Assimes, Absher, Horvath) and by the National Institute of Environmental Health Sciences R01-ES020836 WHI-EMPC (Whitsel, Baccarelli, Hou). The WHI program is funded by the National Heart, Lung, and Blood Institute, National Institutes of Health, U.S. Department of Health and Human Services through contracts HHSN268201100046C, HHSN268201100001C, HHSN268201100002C, HHSN268201100003C, HHSN268201100004C, and HHSN271201100004C. The authors thank the WHI investigators and staff for their dedication, and the study participants for making the program possible. A full listing of WHI investigators can be found at: www.whi.org/researchers/Documents%20%20Write%20a%20Paper/WHI%20Investigator%20Short%20List.pdf.

The InCHIANTI study baseline (1998-2000) was supported as a “targeted project” (ICS110.1/RF97.71) by the Italian Ministry of Health and in part by the U.S. National Institute on Aging (Contracts: 263 MD 9164 and 263 MD 821336).

Funding for the DNA methylation in JHS provided by NHLBI R01HL116446.

The funding bodies played no role in the design, the collection, analysis, or interpretation of the data.

## Availability of data and materials

The WHI data are available at dbGaP under the accession numbers phs000200.v10.p3. The FHS data are available at dbGaP under the accession numbers phs000342 and phs000724. The Normative Aging data are available from dbGAP phs000853.v1.p1. The Jackson Heart Study data are available from https://www.jacksonheartstudy.org/Research/Study-Data

